# Chicken rRNA gene cluster structure

**DOI:** 10.1101/035477

**Authors:** Alexander Demin, Elena Koshel, Alsu Saifitdinova, Svetlana Galkina, Tatsuo Fukagawa, Elena Gaginskaya

**Author notes:** Corresponding author: Elena R. Gaginskaya.

## Abstract

**Background:** Repeated clusters of ribosomal genes whose activity results in the nucleolus formation are extremely important in multicellular organism genome. Despite the extensive exploration into vertebrate genomes, in many model objects the ribosomal cluster structure is still underinvestigated. So far, complete description for primary structure of avian ribosomal cluster has not been reported.

**Results:** This work represents the first successful assembly of complete chicken ribosome cluster sequence. The sequence was deposited to GenBank under accession number KT445934. The total cluster size from *pre-rRNA* transcriptional start site to the 3′ end of *3’ETS* amounted to 11444 bp. *18S rRNA* gene size is 1823 base pairs, *5.8S rRNA –* 157 bp, *28S rRNA –* 4443 bp. The *5 ‘ETS* spacer core size is 1839 bp, *3’ETS –* about 350 bp., *ITS1 –* 2099 bp, and *ITS2 –* 733 bp. The assembly was validated through *in situ* fluorescent hybridization (FISH) analysis on metaphase chromosomes of chicken. *ITS1* and *ITS2* spacer sequences have been found to have high GC pair content and form secondary structures featuring high melting temperature.

**Conclusions:** Decoding of the chicken rRNA gene cluster sequence extends the use of birds as a model object for exploration into nucleolus organizer region (NOR) regulation and nucleolus functions, e.g. in ontogenesis. This data might also be useful to address certain problems of population and evolutionary genetics.

## Background

Repetitive ribosome gene clusters constitute one of the most important genome components, but they are still underinvestigated [1]. They form nucleolus organizing regions (NOR) in chromosomes and the functional status of NOR plays as an indicator of the physiological status of cells, tissues and the entire organism at various ontogenetic stages [2–6]. The precise structure of ribosome cluster contributes to advanced analysis of NOR processes.

All animal ribosome clusters are known to have fundamentally similar structure (Fig. 1). The structure of these clusters is based on sequences of RNA encoding conservative genes *(18S rRNA, 5.8S rRNA* and *28S rRNA)* divided by internal transcribed spacers *(ITS1* and *ITS2)* and flanked by external transcribed spacers *(5’ETS* and *3’ETS)*. All listed elements transcribe into a single RNA predecessor, *pre-rRNA*. Both internal and external transcribed spacers feature high structural variability, which accounts for ribosome cluster length variation within a wide range from 8 to 14 thousand base pairs (bp). The clusters are separated from each other intergenic spacers *(IGS)* containing promoter and terminator regions for RNA Poll that transcribes *pre-rRNA* [7].

**Fig. 1.**
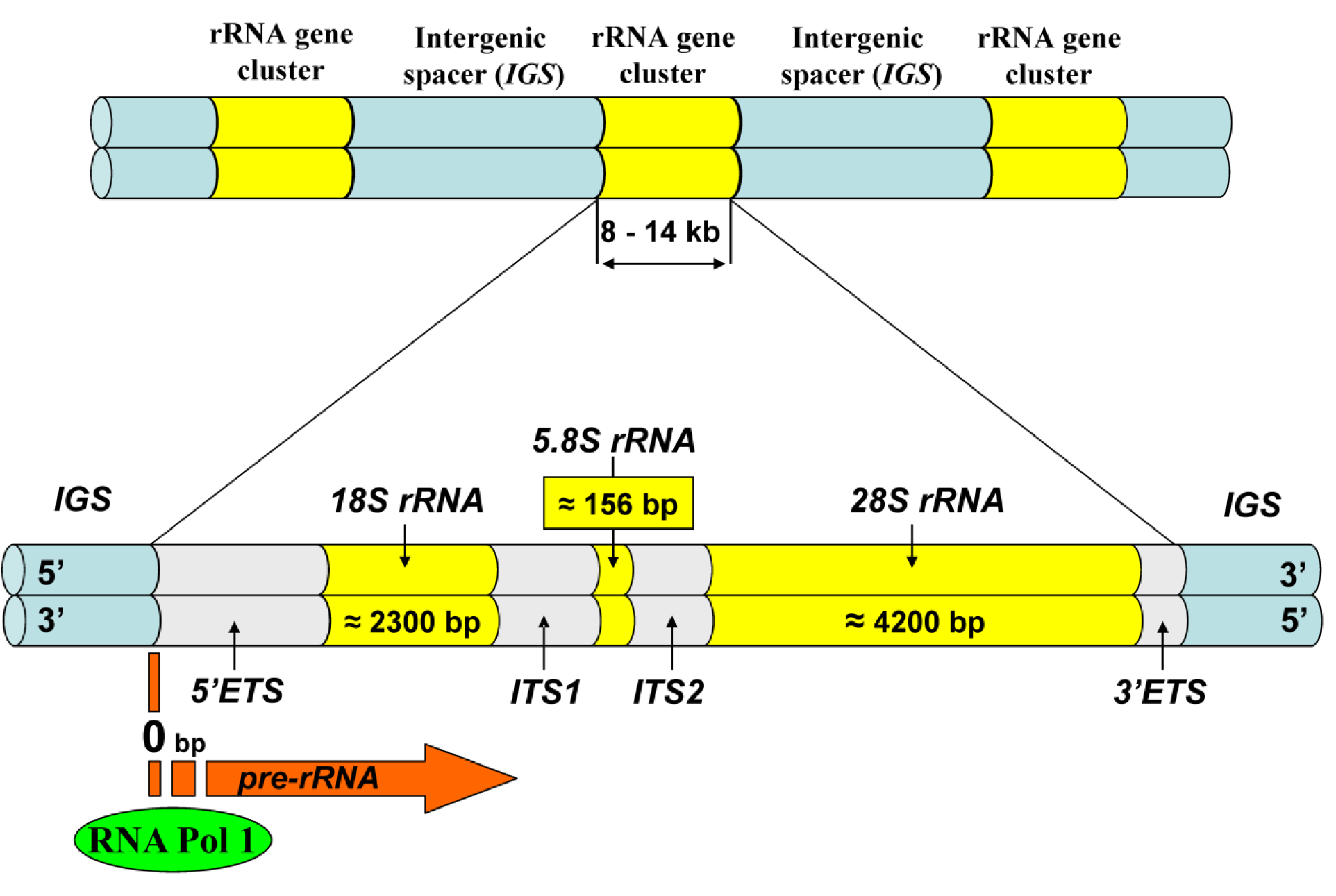
Ribosomal DNA structure (after Singer & Berg, 1991)

Despite extensive studies of vertebrate genomes conducted recently [8], ribosome cluster exploration remains a complicated task, primarily due to high repetitivity and extensive length of the clusters as well as faster spacer evolution [1]. So far GenBank [9] has contained annotated complete rRNA gene cluster sequence only for a limited number of vertebrate species including *Homo sapiens* (GenBank accession number: HSU13369), *Mus musculus* (GenBank accession number: BK000964) and *Nothobranchius furzeri* (GenBank accession number: EU780557). Yet, no description of the complete ribosomal cluster sequence has been offered so far for such a major taxon as Aves.

Chicken ribosomal cluster deserves special research focus among Aves class representatives. Chicken is an important organism for agriculture and in addition, it is well used in extensive range of biomedical research including developmental biology, genetics, cell biology, histology, virology and so on [10, 11]. Chicken genome is firstly sequenced in avian and one of the first among vertebrata genomes [12]. *Gallus gallus 4.0* assembly has been used as a reference genome for assembling sequenced genomes of other birds [13]. Chicken NOR was localized on the chromosome 16 (GGA16) [14–17] and this was recently confirmed by FISH using WAG137G04 BAC clone as a probe [18]. However, the published assembly of Gallus *gallus 4.0* genome [12] has decoded only 5% of GGA16 chromosome sequence and does not include the NOR region [18].

So far, complete description of the primary structure of chicken ribosomal cluster has not been reported. GenBank contains annotated sequences for individual fragments of chicken *18S rRNA* and *28S rRNA* genes. Besides, two groups of authors have contributed to GenBank annotated sequences of *ITS1* and *ITS2* spacers and *5.8S rRNA* gene (Accession number: DQ018752 – DQ018755; FJ008990). However, unfortunately, these sequences are different from each other.

During the course of the “ChIP-sequencing with CENP-A from chicken cells containing neocentromeres on Z chromosome” project (NCBI BioProject accession number: PRJDB2279) [19], we obtained multiple raw chicken sequences, most of which are annotated in Sequence Read Archive (SRA accession numbers: DRX001860 – DRX001863). Based on these sequences as well as sequences of unannotated contigs from the *Gallus gallus 4.0* genome assembly (NCBI WGS accession number: AADN00000000.3) we may clarify a complete structure of chicken ribosomal cluster.

In this study we have assembled and described the complete structure of chicken ribosomal cluster on using an integrated assembly of raw reads available in SRA, WGS contigs and Nucleotide database sequences (GenBank). Finally, we validated our analysis by FISH (fluorescent hybridization) on mitotic chromosomes. The results of this study may contribute notably to expansion of avian use as a model object, in particular, for exploring NOR regulation in ontogenesis. The output data would be useful genetics and evolutional biology.

## Methods

### Cluster assembling based on published data

To assemble a complete chicken ribosomal cluster we used the sequence library [19] which had been earlier generated of raw reads (SRA accession number: DRX001863) sequenced in the course of the “ChIP-sequencing with CENP-A from chicken cells containing neocentromeres on Z chromosome” project (NCBI BioProject accession number: PRJDB2279). For assembly verification and rectification we used 19 unannotated sequences of the Whole Genome Shotgun (WGS) contigs from the *Gallus gallus 4.0* genome assembly (NCBI WGS accession number: AADN00000000) [12] and 9 sequences from Nucleotide database [20] annotated by the authors as chicken ribosomal cluster fragments (Table 1).

The search for raw sequences and WGS contigs homologous to the ribosomal cluster elements was performed using BLAST [22]. For raw read tiling, alignment of contigs and annotated sequences and nucleotide structure determination, UGENE 1.16.1. [23] and Mega 6.06 [24] were used. Repeat search and typing was performed in Repeatmasker 4.0.5 [25]. Nucleotide sequence secondary structure was recreated in Mfold [26] software.

**Table 1.** WGS contigs and annotated sequences used to verify the chicken ribosomal cluster assembly

| Cluster element | Accession number | Reference |
| --- | --- | --- |
| 5'ETS<br>(complete) | NW_003775878, AADN03015064<br>AADN03001778, AADN03014081<br>AADN03001785, AADN03001786<br>AADN03001788<br>DQ112354 | [12]<br>[11] |
| 18S rRNA<br>(complete) | AADN03000430, AADN03001677<br>AADN03001785, AADN03001786<br>AADN03001788, AADN03026634<br>FM165414<br>DQ018752, DQ018754 | [12]<br>Unpublished<br>Unpublished |
| ITS1<br>(complete) | AADN03026634, AADN03000430<br>AADN03001782, AADN03001670<br>AADN03001783<br>DQ018752, DQ018754 | [12]<br>Unpublished |
| 5.8S rRNA<br>(complete) | AADN03001670, AADN03001783<br>DQ018754 | [12]<br>Unpublished |
| ITS2<br>(complete) | AADN03001783, AADN03001784 | [12] |
| 28S rRNA<br>(complete) | AADN03001784, AADN03001774<br>AADN03001775, AADN03001776<br>AADN03022685, AADN03019346<br>EF552813<br>FM165415<br>JN639848<br>DQ018756, DQ018757 | [12]<br>[21]<br>Unpublished<br>Unpublished<br>Unpublished |

To define the boundaries between spacer *(5’ETS, ITS1, ITS2, 3’ETS)* sequences and *rRNA (18S, 5.8S, 28S)* gene sequences we used annotated fragments of ribosomal clusters of *Homo sapiens* (GenBank accession number: HSU13369), *Rattus norvegicus* (GenBank accession number: NR_046239), *Mus musculus* (GenBank accession number: NR_046233), *Xenopus laevis* (GenBank accession number: X02995) and *Crocodylus porosus* (GenBank accession number: EU727191).

### Experimental confirmation of assembly accuracy

To confirm the accuracy of our assembly and the location of the assembled sequence to the NOR on chromosome GGA16, serial fluorescent hybridization *in situ* (serial FISH) of assembled sequence fragments and WAG137G04 BAC clone known to include a chicken GGA16 fragment comprising NOR [18], was applied to chicken mitotic chromosomes.

The mitotic chromosomes were obtained from fibroblasts of a four-day chicken embryo by standard procedure.

Probes to the assembled sequence were PCR amplified boundary areas of *5’ETS–18S rRNA* and *ITS1–5.8S rRNA*. PCR primers were designed based on the assembled rRNA cluster (Table 2) using Unipro UGENE 1.16.1 software; their identity to the related regions of the assembled rRNA gene cluster was validated by standard sequencing. WAG137G04 BAC clone probe was produced by standard DOP-PCR amplification using 6MW primers [27].

PCR amplification of *5’ETS-18S rRNA* and *ITS1-5.8S rRNA* areas was carried out in 20 ml of reaction mix as follows: Taq-pol 5U/μl (Sileks) – 0,5 μl; 10X Taq Buffer (Sileks) – 2 μl; MgCl2 25 mM (Sileks) – 2 μl; 10mM dNTP (2.5 mM each) (Sileks) – 1.6 μl; 10μM primer (10 pmol/μl) – пo 1 μl each; DNA – 0,5 μl H_2_O – 11.4 μl.

PCR protocol was as follows: 94°C–5’; (94°C–20”, 60°C–15”, 72°C–20”)x35n; 72°C–5’; 4°C–hold.

**Table 2.**
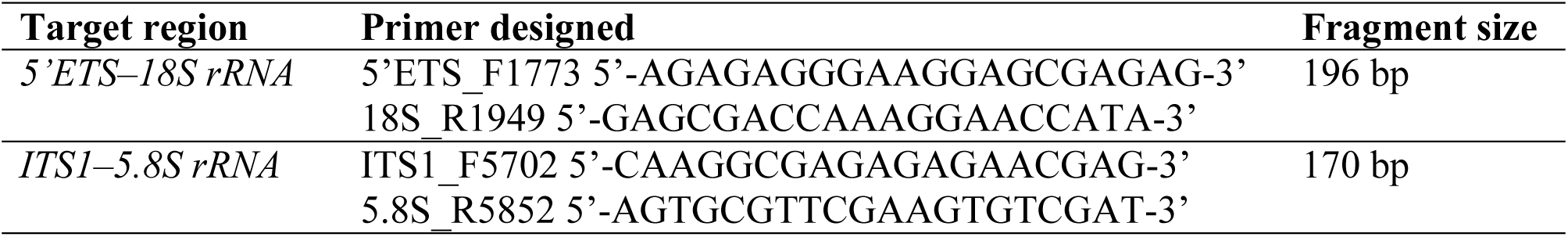
The list of primers for amplification of target regions of the assembled ribosomal cluster sequence

Sequencing of PCR products was carried out at the Molecular and Cell Technology Development Saint-Petersburg State University Resource Center.

Fluorescent probes were generated by labeling PCR products with modified biotin-16-dUTP nucleotide (Sileks) during amplification procedure.

FISH was carried out according to a published protocol [28]. The preps were additionally stained with 4’,6-diamidino-2-phenilindole-dihydrochloride (DAPI) in concentration of 1μg/ml.

For establishing whether the fluorescent probe hybridization had taken place at NOR on GGA16, re-FISH with *WAG137G04* BAC clone probe was carried out on the same preparations.

FISH results were investigated using DM4000B (Leica) epifluorescent microscope in the Chromas Saint-Petersburg Recourse Center. The images were processed and superposed using Adobe Photoshop CS5.1 software.

### Results and discussion

The assembled and annotated cluster is 11444 bp in size and contains complete *5’ETS*, *18S rRNA*, *5.8S rRNA*, *ITS2*, *28S rRNA* sequences and 5’-end of *3’ETS* sequence. This cluster sequence was deposited to GenBank under accession number KT445934.

### rRNA gene structure

#### 18S rRNA

We used the raw sequences deposited to SRA under accession number DRX001863 to assemble the complete sequence of *18S rRNA* gene. To confirm accuracy of our assembly, six WGS contigs and three annotated sequences of fragments of this gene were used in addition to raw sequences (Table 1). These WGS contigs and annotated sequences covered 100% and 99.4% of the assembly correspondingly (Additional file 1). Within the designated boundaries (Fig. 2) the size of the complete *18S rRNA* sequences was 1823 bp (from 1835 through 3662 bp starting from the transcription initiation site; Additional file 2), which is virtually the same as the size of *Xenopus laevis 18S rRNA* (1825 bp). GC pair content is 54.5% and is also comparable to other taxon data: *Homo sapiens* (56.1%), *Xenopus laevis* (53.8%), *Crocodylus porosus* (49.5%).

**Fig. 2.**
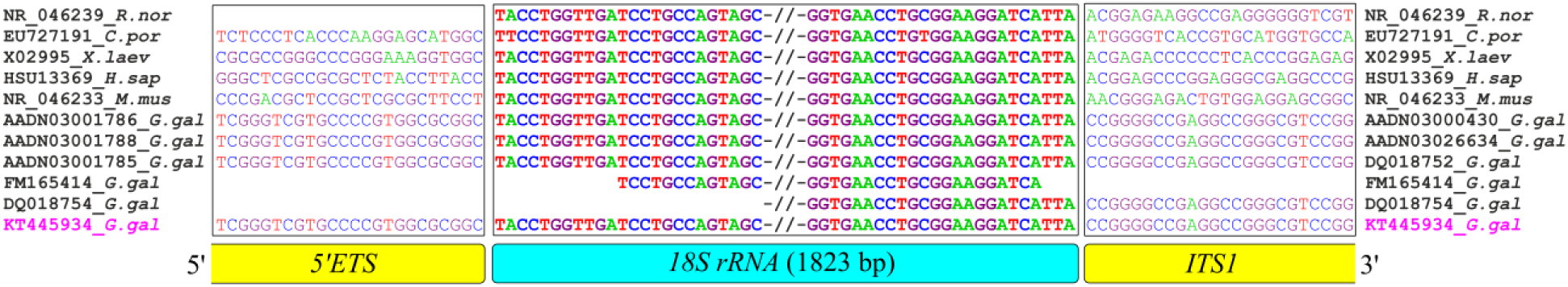
*18S rRNA* gene sequence boundaries resulted from a multiple sequence alignment. Black frames outline the elements of the ribosomal cluster. Cluster members are schematically indicated by colored boxes. Double slashes schematically point the hidden area of alignment. Extreme right and left – accession numbers and taxa abbreviations relating to the sequences located before and after the hidden area of the alignment, respectively. G. gal – Gallus gallus; R. nor – *Rattus norvegicus*; C. por – *Crocodylus porosus*; X. laev – *Xenopus laevis;* H. sap – *Homo sapiens*; M. mus – *Mus musculus*. KT445934_G.gal – sequence assembled by raw read tiling

#### 5.8S rRNA

The complete sequence of chicken *5.8S rRNA* gene was assembled on the basis of raw reads. To validate this assembly two WGS contigs and annotated sequence (GenBank accession number: DQ018754) (Table 1) were used. These WGS contigs and annotated sequences covered 100% and 66.2% of the assembly respectively (Additional file 1). Upon alignment (Fig. 3) the size of chicken *5.8S rRNA* gene was 157 bp (from 5762 through 5918 bp) and similar to the equivalent human gene (Additional file 2). The number of GC pars in chicken *5.8S rRNA* sequence was also similar to human (57.3% and 57.4% respectively).

**Fig. 3.**
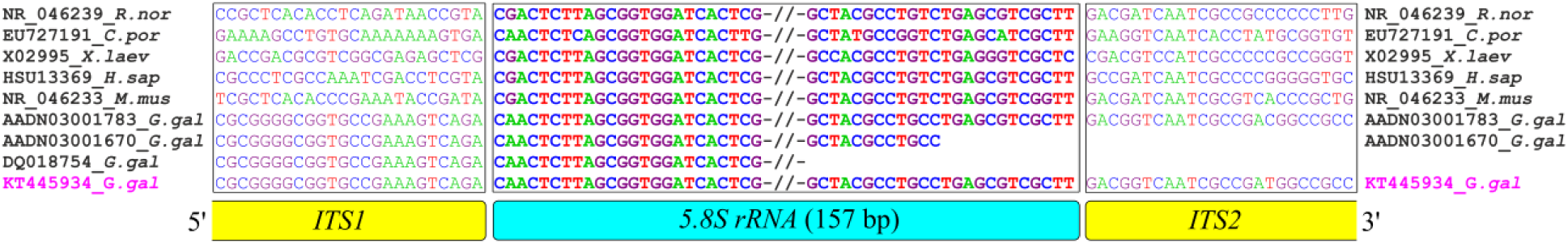
Chicken *5.8S rRNA* gene sequence boundaries resulted from a multiple sequence alignment. Black frames outline the elements of the ribosomal cluster. Cluster members are schematically indicated by colored boxes. Double slashes schematically point the hidden area of alignment. Extreme right and left – accession numbers and taxa abbreviations relating to the sequences located before and after the hidden area of the alignment, respectively. G. gal – Gallus gallus; R. nor – *Rattus norvegicus*; C. por – *Crocodylus porosus*; X. laev – *Xenopus laevis*; H. sap – *Homo sapiens*; M. mus – *Mus musculus*. KT445934_G.gal – sequence assembled by raw read tiling

#### 28S rRNA

The structure of chicken *28S rRNA* sequence was assembled using raw sequences from SRA and involved six WGS contigs, five annotated sequences from GenBank and *28S rRNA* gene sequence (Table 1). These WGS contigs and annotated sequences covered 76.7% and 69.7% of the assembly respectively (Additional file 1). Within the designated boundaries (Fig. 4) chicken *28S rRNA* gene size is 4443 bp (from 6652 through 11094 bp) (Additional file 2). GC pair content in *28S rRNA* chicken gene is much higher than in *18S* and *5.8S rRNA* genes and achieves as much as 68.0%. *28S rRNA* gene sequence features a heterogeneous structure: it contains 2 continuous fragments with increased GC pair content (>70% in average): from 7115 through 8067 bp and from 9301 through 9746 bp.

**Fig. 4.**
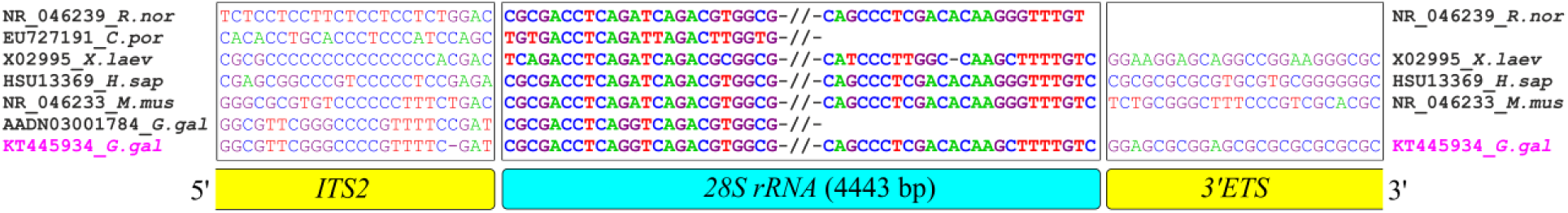
Chicken *28S rRNA* gene sequence boundaries resulted from a multiple sequence alignment. Black frames outline the elements of the ribosomal cluster. Cluster members are schematically indicated by colored boxes. Double slashes schematically point the hidden area of alignment. Extreme right and left – accession numbers and taxa abbreviations relating to the sequences located before and after the hidden area of the alignment, respectively. G. gal – Gallus gallus; R. nor – *Rattus norvegicus*; C. por – *Crocodylus porosus*; X. laev – *Xenopus laevis;* H. sap – *Homo sapiens*; M. mus – *Mus musculus*. KT445934_G.gal – sequence assembled by raw read tiling

### Internal transcribed spacers *(ITS)*

#### ITS1

*ITS1* sequence was deposited to GenBank by Tang et al. under accession numbers DQ018754 (complete sequence) and DQ018752 (partial sequence), and by Chen et al. under accession number FJ008990 (complete sequence). These sequences do not match each other. We tried to clarify chicken *ITS1* structure using tiling assembly of raw sequences and 5 WGS contig data. The WGS contigs covered 100% of the assembly (Additional file 1). Within the designated boundaries (Fig. 5) the size of chicken *1TS1* was 2099 bp (from 3663 through 5761 bp) (Additional file 2). Our whole sequences for *ITS1* are homologous to the version offered by Tang et al., with the exception of spacer boundaries. Tang et al. report its size as 2155 bp due to inclusion of spacer flank sequences of *18S rRNA* and the entire *5.8S rRNA. ITS1* nucleotide content analysis has resulted in high GC pair content – 81.4% and high CpG dinucleotide content – 19.7%. Repeats account for 18.2% of *ITS1* size and are composed of the following nucleotide combinations: (CCGAGG)n, (CCGGT)n, (GC)n, (GGGGGCC)n, (GAG)n и (GGCGCG)n. High GC pair content in *ITS1* sequence leads to formation of secondary structures with multiple hairpins. In accordance with the models produced for T – 60°C, [K^+^] – 50 mM PCR conditions, complete hairpin dissociation does not take place (Additional file 3). Under such conditions, the average increment of Gibbs free energy (ΔG) in ITS1 sequence in single-stranded DNA form is -130.63.

**Fig. 5.**
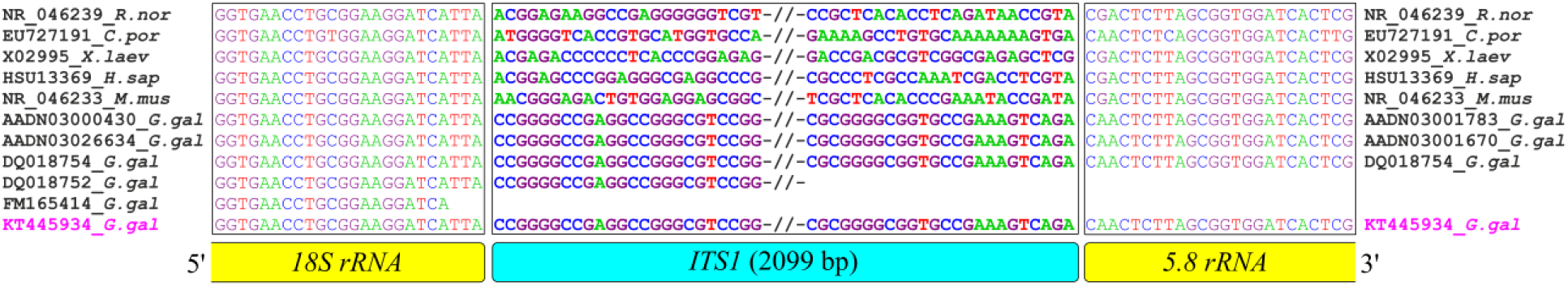
Chicken *ITS1* sequence boundaries resulted from a multiple sequence alignment. Black frames outline the elements of the ribosomal cluster. Cluster members are schematically indicated by colored boxes. Double slashes schematically point the hidden area of alignment. Extreme right and left – accession numbers and taxa abbreviations relating to the sequences located before and after the hidden area of the alignment, respectively. G. gal – Gallus gallus; R. nor – *Rattus norvegicus*; C. por – *Crocodylus porosus*; X. laev – *Xenopus laevis*; H. sap – *Homo sapiens*; M. mus – *Mus musculus*. KT445934_G.gal – sequence assembled by raw read tiling

#### ITS2

Chicken *ITS2* assembly from raw sequences was validated using two overlapping WGS contigs (Table 1), assembly coverage being 100% (Additional file 1). We excluded from the sequences deposited in GenBank with accession numbers DQ018753, DQ018755 and FJ008990, which are annotated as containing chicken *ITS2*. The reason was that the related gene flank sequences were found non-homologous to 3’-end of *5.8S rRNA* sequence and to *5*’-end of chicken *28S rRNA*. Within the designated boundaries (Fig. 6) the size of chicken *1TS2* was 733 bp (from 5919 through 6651 bp) (Additional file 2). Repeats account for 24.3% of spacer size. These repeats are composed of the following nucleotide combinations: (CCGT)n, (GCGCG)n, (CGTT)n, (CCGT)n, (GCG)n. The sequence features increased GC pair content – 82.0% and CpG dinucleotide content – 20.7% which is virtually similar to *ITS1* values (see above). Similarly to *ITS1, ITS2* also feature formation of extended hairpins with high melting temperature. According to our results obtained under standard PCR parameters, ΔG = −62.5 (Additional file 4).

**Fig. 6.**
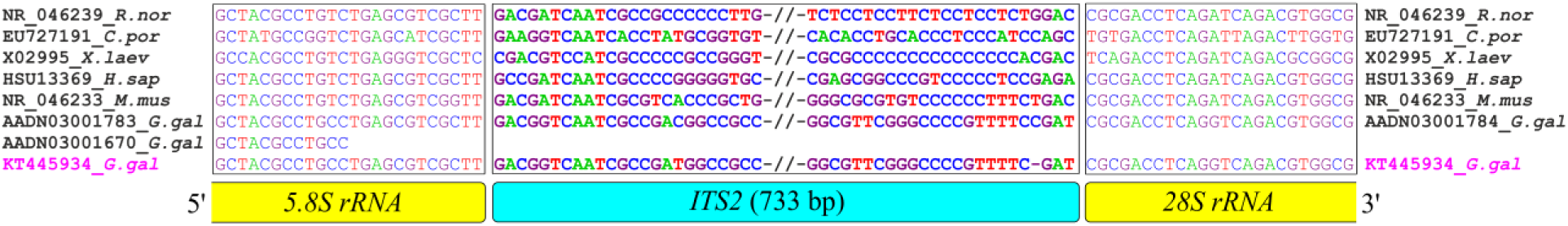
Chicken *ITS2* sequence boundaries resulted from a multiple sequence alignment. Black frames outline the elements of the ribosomal cluster. Cluster members are schematically indicated by colored boxes. Double slashes schematically point the hidden area of alignment. Extreme right and left – accession numbers and taxa abbreviations relating to the sequences located before and after the hidden area of the alignment, respectively. G. gal – Gallus gallus; R. nor – *Rattus norvegicus*; C. por – *Crocodylus porosus*; X. laev – *Xenopus laevis*; H. sap – *Homo sapiens*; M. mus – *Mus musculus*. KT445934_G.gal – sequence assembled by raw read tiling

Chicken *ITS1* sequence has proved to be more extended than that of most animals, with the exception of marsupials [29]. Chicken *ITS2* has also proved to be far more extended than it had been reported previously (GenBank accession numbers: DQ018753, DQ018755, FJ008990). Our attempts to amplify and obtain complete chicken *ITS1* and *ITS2* sequences using traditional approaches have been unsuccessful. Our findings suggest that the key problem in avian *ITS1* and *ITS2* amplification and sequencing may probably be related to high CG pair content and secondary structure formation. These factors impact polymerase effect in PCR process and increase the probability of AT-enriched regions non-specific amplification. This factor is quite likely to be the main reason for the current unavailability of annotated extended avian ribosomal cluster sequences in GenBank. Representativity of animal *ITS1* and *ITS2* sequences in GenBank is shown on the Table 3.

Notably, among the eight avian sequences deposited to GenBank, only three actually belong to the ribosomal cluster, namely to *ITS1* sequence. At the same time, complete deciphering of the ribosomal cluster sequence in different groups of organisms is of a great value for various fields of biology, primarily systematics, phylogeny and ecology. Our alignment of chicken rRNA gene complete cluster might be useful for comparative research in these fields. In the majority of animals, including birds, the order of alternation of coding and spacer sequences within rRNA gene clusters is conserved. Yet spacer sequences, particularly *ITS1* and *ITS2*, are characterized by high rates of variability. Due to this they are used extensively as DNA barcodes as well as nuclear markers of micro- and macro-evolutonary events [30–33]. However, the features of avian *ITS1* and *ITS2* do not allow treating their sequences as easily accessible and effective DNA barcodes or phylogenetic markers for this class.

**Table 3.** GenBank representation of *ITS1* and *ITS2* sequences

| Taxonomic group | Deposited sequences |  |
| --- | --- | --- |
|  | *ITS1* | *ITS2* |
| Nematoda | 7301 | 7283 |
| Insecta | 9616 | 19614 |
| Mollusca | 5729 | 4374 |
| Chondrichthyes | 22 | 193 |
| Osteichthyes | 1432 | 571 |
| Amphibia | 4 | 51 |
| Reptilia | 56 | 102 |
| **Aves** | **5** | **3** |
| Mammalia | 22 | 23 |

### External transcribed spacers *(ETS)*

#### 5’ETS

The validity of our chicken rRNA *5’ETS* assembly was confirmed using seven WGS contigs (Table 1). Additionally, we used a chicken sequence (GenBank accession number: DQ112354) containing the entire RNA PolI promoter and a *pre-rRNA* initiator site [11]. These WGS contigs and annotated sequences covered 63.9% and 14.8% of the assembly respectively (Additional file 1). Within the designated boundaries (Fig. 7) the size of *5’ETS* is 1839 bp (from 1 through 1832 bp) (Additional file 2). Within the assembled *5 ‘ETS*, repeating sequences account for 6.4% of its size and are composed of (GCGA)n and (GAGA)n nucleotide combinations and (CGG)n and (GCC)n reverse repeats, the latter being responsible for formation of hairpins related to rRNA processing. The *5 ‘ETS* sequence was also found to feature increased GC pair content – 75.0%. At the same time, CpG content in chicken *5’ETS* (15.7%) is comparable to CpG content in 5’ETS of human (16.6%) and Xenopus laevis (15.5%).

**Fig. 7.**
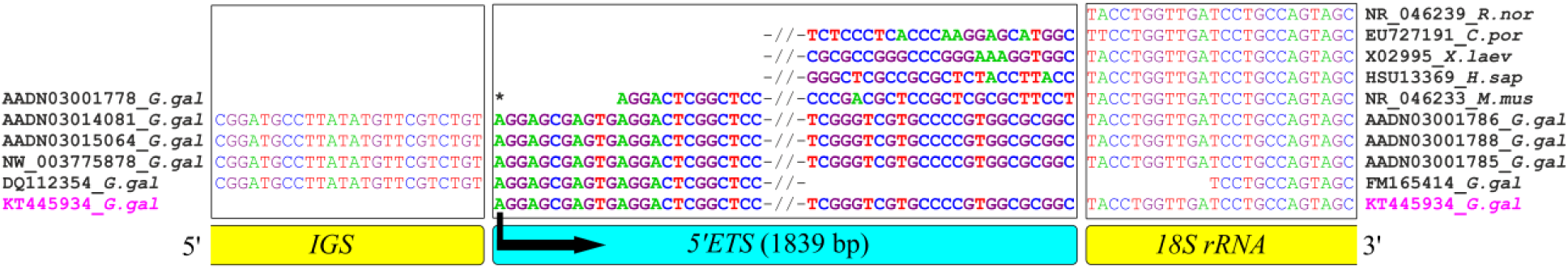
Chicken *5’ETS* sequence boundaries resulted from a multiple sequence alignment. Black frames outline the elements of the ribosomal cluster. Cluster members are schematically indicated by colored boxes. Asterisk – chicken *pre-rRNA* transcription initiation site. Arrow – transcription direction. Double slashes schematically point the hidden area of alignment. Extreme right and left – accession numbers and taxa abbreviations relating to the sequences located before and after the hidden area of the alignment, respectively. G. gal – Gallus gallus; R. nor – *Rattus norvegicus;* C. por – *Crocodylus porosus;* X. laev – *Xenopus laevis;* H. sap – *Homo sapiens;* M. mus – *Mus musculus*. KT445934_G.gal – sequence assembled by raw read tiling

Despite significant differences in 5’ETS length among these three species (1961, 3656 and 713 bp, respectively) and lack of a clear homology (Additional file 2), their CpG content variability does not exceed 1.1%. The CpG dinucleotide content similarity in varying sequences from quite distant animal taxa probably suggests the importance of CpG role in promoter and spacer functions of *5’ETS*. It might also be indicative of the existence of a stabilizer of spacer nucleotide content.

#### 3’ETS

Chicken *3’ETS* sequence was assembled from raw reads. The size of the initial assembly was 481 bp. Yet we were unable to define precisely the position of the 3’-end of *3’ETS* due to lack of data on the sequence of Sal1-box, the variable element of termination of the transcription by RNA-Pol 1 [34].

Based on comparison of sizes and nucleotide content of *3’ETS* in other organisms (345 bp/82.9%GC in human, 521 bp/75.62% GC in mouse, 236 bp/84.32% GC in frog), it is possible to assume that termination of *pre-rRNA* transcription in chicken could occur at 11444 bp position. In chicken *3’ETS* sequence the region between 11445 bp and 11462 bp is a poly-T region (Additional file 1). This region has not been reported to exist in *3 ‘ETS* of *Homo sapiens, Mus musculus* and *Xenopus laevis* (Additional file 5). An analysis of 350 bp (from 11095 through 11444 bp) of the *3 ‘ETS* region has resulted in 80.3% and 18.3% of GC pair and CpG dinucleotide content respectively. (GC)n, (CGTT)n and (CGGC)n repeats constitute about 30.9% of the analyzed sequence. These results indicate a certain similarity between the *3 ‘ETS* and other the ribosomal cluster spacers. We believe that it could be related to the existence of a general evolutionary mechanism supporting this stability of spacer nucleotide content within the ribosomal cluster sequence in birds.

### Fluorescent *in situ* hybridization

To verify the relation of the assembled sequence to chicken ribosomal cluster and its NOR location on GGA16, we applied serial re-FISH to chicken mitotic chromosomes. Probes for boundary areas of *5’ETS-18S rRNA* and *ITS1-5.8S rRNA* within the assembled sequence were obtained from genome DNA by primer synthesis. Their sizes were 196 bp and 170 bp respectively. The primers were calculated based on the assembled rRNA cluster. Probe identity to the related regions of the assembled rDNA cluster was validated by standard sequencing. The NOR detection probe was produced by DOP-PCR amplification of BAC-clone WAG137G04 known to include a chicken GGA16 fragment comprising NOR [18]. On chicken mitotic plates, both the spacer and NOR probes hybridized on the same sites in two microchromosomes (Fig. 8). Our findings strongly confirm the reliability of our assembly of chicken ribosomal cluster.

**Fig. 8.**
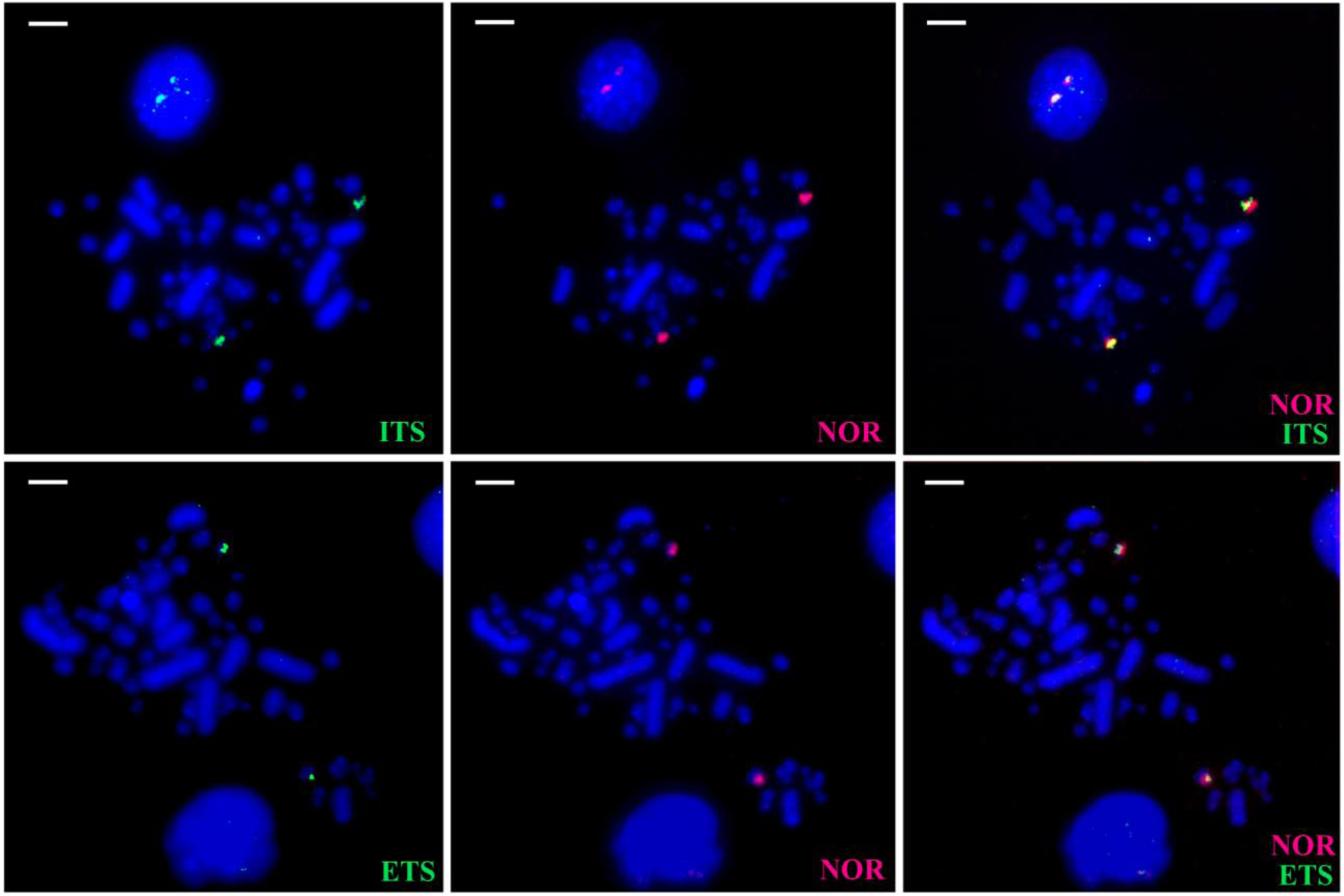
Fluorescent probe co-localization with NOR on mitotic GGA16. ETS – green fluorescent signal of *5’ETS-18S rRNA* boundary fragment probe; ITS – green fluorescent signal of *IT1S-5.8S rRNA* boundary fragment probe; NOR – red fluorescent signal of *WAG137G04* BAC clone probe. Bar – 5 μ

## Conclusion

In this work, we have determined, verified, and featured the complete sequence of chicken ribosomal cluster (Fig. 9). Codings of *18S, 5.8S* and *28S rRNA* gene sequences have typical for higher vertebrate structures. Both *ITS1* and *ITS2* were found to be of a longer size and GC higher content. As a result, they have a complicated secondary structure preventing their PCR analysis and consequently their use as phylogenetic markers. It also makes chicken ribosomal genome analysis complicated in total. It seems more promising to use a relatively less GC-enriched *5’ETS* sequence for the above purpose. Meanwhile, *ITS1, ITS2* and *3’ETS* sequences revealed similarities in the GC, CpG and repeated sequence contents. Thus it is possible to suggest the existence of a general evolutionary mechanism supporting the spacer constant nucleotide proportions within avian rDNA genome.

**Fig. 9.**
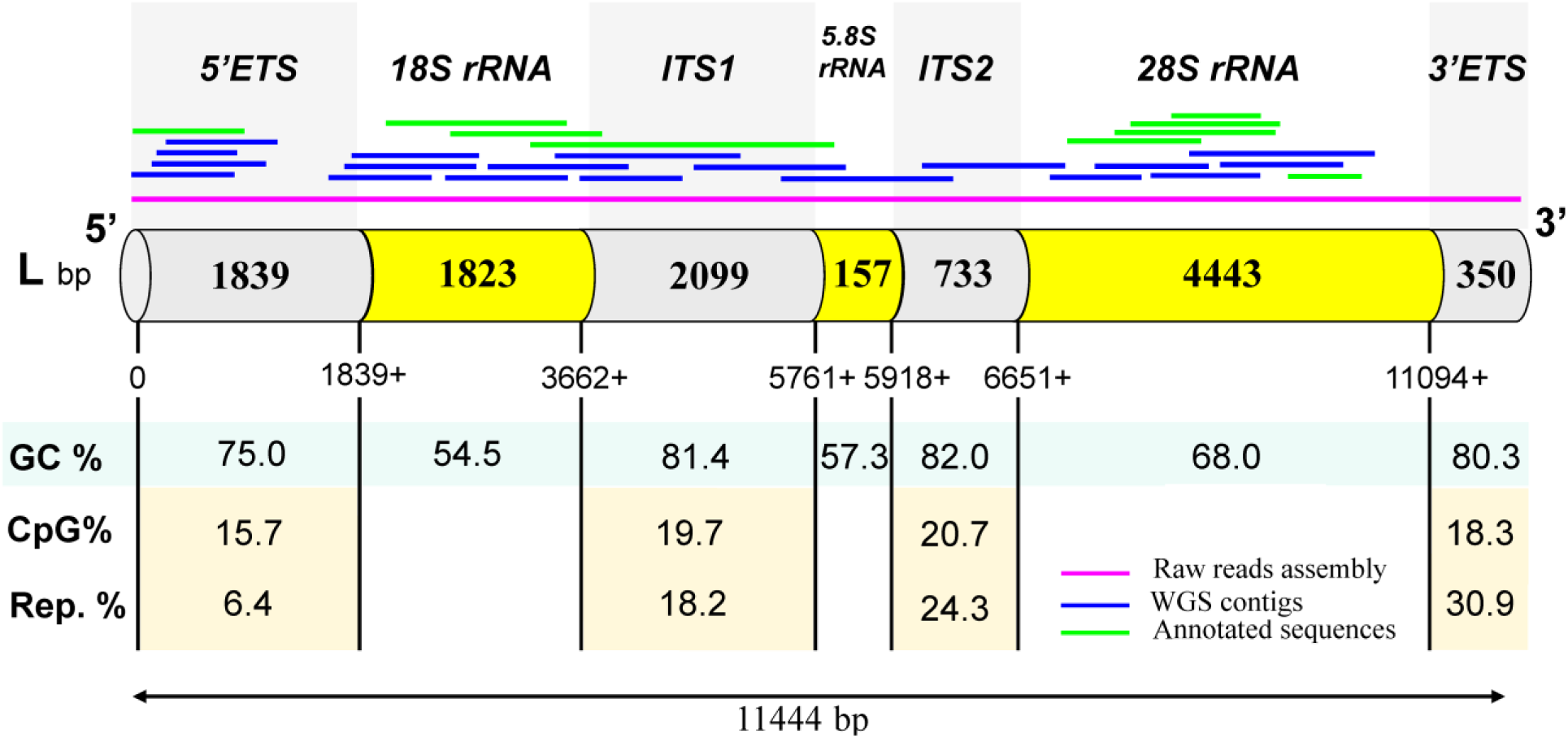
The chicken rRNA gene cluster structure and features. They were established on the basis of data from three genetic databases: raw reads assembly (SRA accession number: DRX001863), WGS contigs (WGS: AADN00000000.3) and early annotated sequences (Nucleotide database)

Knowledge of the chicken rRNA gene cluster structure extends the use of birds as a model object for exploration of the NOR regulation, particularly in ontogenesis and cell differentiation. The results obtained can be useful for searching optimal solutions in the areas of population genetics and evolution.

## Availability of supporting data

The data sets supporting the results of this article are included within the article and its additional files.

**Abbreviations**
SRA: – Sequence Read Archive
WGS: – Whole Genome Shotgun
NOR: – Nucleolus Organizer Regions
5’ETS: *– 5’ External Transcribed Spacer*
3’ETS: *– 3’ External Transcribed Spacer*
ITS1: *– Internal Transcribed Spacer 1*
ITS2: *– Internal Transcribed Spacer 2*
PCR: – Polymerase Chain Reaction
BAC: – Bacterial artificial chromosome
FISH: – Fluorescence In Situ Hybridization

## Competing interests

The authors declare that they have no competing interests.

## Authors’ contributions

AD – chicken ribosomal cluster sequence assembling; EK and SG – metaphase chromosome preparing and FISH; TF – chicken ribosomal cluster fragment sequencing; AS and EG – participation in the study design and coordination. All authors critically revised the manuscript and gave approval of the final version.

## Acknowledgements

The research has been financed by Saint-Petersburg State University (project number 1.50.1043.2014) and Russian Basic Research Foundation (project 15-04-05684). Saint-Petersburg State University Scientific Park’s resources were used.

## Additional files

**Additional file 1:** Raw read assembly verification using WGS contigs and annotated sequences.

**Additional file 2:** Multiple alignment matrix for chicken rRNA gene cluster boundaries search using annotated fragments of ribosomal clusters of *Homo sapiens, Rattus norvegicus, Mus musculus, Xenopus laevis* and *Crocodylus porosus*.

**Additional file 3:** Chicken *ITS1* ssDNA secondary structures under the conditions of T = 60°C, [K^+^] = 50 mM.

**Additional file 4:** Chicken *ITS2* ssDNA secondary structures under the conditions of T = 60°C, [K^+^] = 50 mM.

**Additional file 5:** The comparison of *3’ETS* sequences in chicken and other vertebrates.

## References

1. Shaw P, Brown J. Nucleoli: composition, function, and dynamics. Plant Physiol. 2012; 158(1):44–51.

2. Smirnov E, Kalmárová M, Koberna K, Zemanová Z, Malínský J, Masata M, Cvacková Z, Michalová K, Raska I. NORs and their transcription competence during the cell cycle. Folia Biol. 2006;52(3):59–70.

3. Hernandez-Verdun D, Roussel P, Thiry M, Sirri V, Lafontaine DL. The nucleolus: structure/function relationship in RNA metabolism. Wiley Interdiscip Rev RNA. 2010;1(3):415–31.

4. Bozinovic G, Sit TL, Hinton DE, Oleksiak MF. Gene expression throughout a vertebrate’s embryogenesis. BMC Genomics. 2011;doi: 10.1186/1471-2164-12-132.

5. Zheng Z, Jia JL, Bou G, Hu LL, Wang ZD, Shen XH, Shan ZY, Shen JL, Liu ZH, Lei L. rRNA genes are not fully activated in mouse somatic cell nuclear transfer embryos. J Biol Chem. 2012;287(24):19949–60.

6. Tafforeau L, Zorbas C, Langhendries JL, Mullineux ST, Stamatopoulou V, Mullier R, Wacheul L, Lafontaine DL. The complexity of human ribosome biogenesis revealed by systematic nucleolar screening of Pre-rRNA processing factors. Mol Cell. 2013;51(4):539–51.

7. Singer M, Berg P. Genes and Genomes, a Changing Perspective. Mill Valley: University Science Books; 1991.

8. Koepfli KP, Paten B. Genome 10K Community of Scientists, O’Brien SJ. The Genome 10K Project: a way forward. Annu Rev Anim Biosci. 2015;3:57–111.

9. NCBI GenBank database. http://www.ncbi.nlm.nih.gov/genbank. Accessed 20 Aug 2015.

10. Zhang X, Yelbuz TM, Cofer GP, Choma MA, Kirby ML, Johnson GA. Improved preparation of chick embryonic samples for magnetic resonance microscopy. Magn Reson Med. 2003;49(6):1192–5.

11. Massin P, Rodrigues P, Marasescu M, van der Werf S, Naffakh N. Cloning of the chicken RNA polymerase I promoter and use for reverse genetics of influenza A viruses in avian cells. J Virol. 2005;79(21): 13811–6.

12. International Chicken Genome Sequencing Consortium. Sequence and comparative analysis of the chicken genome provide unique perspectives on vertebrate evolution. Nature. 2004;432(7018):695–716.

13. Zhang G, Li C, Li Q, Li B, Larkin DM, Lee C, Storz JF, Antunes A, Greenwold MJ, Meredith RW, Ödeen A, Cui J, Zhou Q, Xu L, Pan H, Wang Z, Jin L, Zhang P, Hu H, Yang W, Hu J, Xiao J, Yang Z, Liu Y, Xie Q, Yu H, Lian J, Wen P, Zhang F, Li H, Zeng Y, Xiong Z, Liu S, Zhou L, Huang Z, An N, Wang J, Zheng Q, Xiong Y, Wang G, Wang B, Wang J, Fan Y, da Fonseca RR, Alfaro-Núñez A, Schubert M, Orlando L, Mourier T, Howard JT, Ganapathy G, Pfenning A, Whitney O, Rivas MV, Hara E, Smith J, Farré M, Narayan J, Slavov G, Romanov MN, Borges R, Machado JP, Khan I, Springer MS, Gatesy J, Hoffmann FG, Opazo JC, Håstad O, Sawyer RH, Kim H, Kim KW, Kim HJ, Cho S, Li N, Huang Y, Bruford MW, Zhan X, Dixon A, Bertelsen MF, Derryberry E, Warren W, Wilson RK, Li S, Ray DA, Green RE, O’Brien SJ, Griffin D, Johnson WE, Haussler D, Ryder OA, Willerslev E, Graves GR, Alström P, Fjeldså J, Mindell DP, Edwards SV, Braun EL, Rahbek C, Burt DW, Houde P, Zhang Y, Yang H, Wang J; Avian Genome Consortium, Jarvis ED, Gilbert MT, Wang J. Comparative genomics reveals insights into avian genome evolution and adaptation. Science. 2014;346(6215):1311–20.

14. Auer H, Mayr B, Lambrou M, Schleger W. An extended chicken karyotype, including the NOR chromosome. Cytogenet Cell Genet. 1987;45(3–4):218–21.

15. Miller MM, Goto RM, Taylor RL Jr, Zoorob R, Auffray C, Briles RW, Briles WE, Bloom SE. Assignment of Rfp-Y to the chicken major histocompatibility complex/NOR microchromosome and evidence for high-frequency recombination associated with the nucleolar organizer region. Proc Natl Acad Sci USA. 1996;93(9):3958–62.

16. Wain HM, Toye AA, Hughes S, Bumstead N. Targeting of marker loci to chicken chromosome 16 by representational difference analysis. Anim Genet. 1998;29(6):446–52.

17. Masabanda JS, Burt DW, O’Brien PC, Vignal A, Fillon V, Walsh PS, Cox H, Tempest HG, Smith J, Habermann F, Schmid M, Matsuda Y, Ferguson-Smith MA, Crooijmans RP, Groenen MA, Griffin DK. Molecular cytogenetic definition of the chicken genome: the first complete avian karyotype. Genetics. 2004;166(3):1367–73.

18. Solinhac R, Leroux S, Galkina S, Chazara O, Feve K, Vignoles F, Morisson M, Derjusheva S, Bed’hom B, Vignal A, Fillon V, Pitel F. Integrative mapping analysis of chicken microchromosome 16 organization. BMC Genomics. 2010;doi: 10.1186/1471-2164-11-616.

19. Shang WH, Hori T, Martins NM, Toyoda A, Misu S, Monma N, Hiratani I, Maeshima K, Ikeo K, Fujiyama A, Kimura H, Earnshaw WC, Fukagawa T. Chromosome engineering allows the efficient isolation of vertebrate neocentromeres. Dev Cell. 2013;24(6):635–48.

20. NCBI Nucleotide database. http://www.ncbi.nlm.nih.gov/nuccore. Accessed 20 Aug 2015.

21. Paśko Ł, Ericson PG, Elzanowski A. Phylogenetic utility and evolution of indels: a study in neognathous birds. Mol Phylogenet Evol. 2011;61(3):760–71.

22. NCBI BLAST databases. http://blast.ncbi.nlm.nih.gov/Blast.cgi. Accessed 15 Aug 2015.

23. Unipro UGENE. http://ugene.net. Accessed 15 Aug 2015.

24. Tamura K, Stecher G, Peterson D, Filipski A, Kumar S. MEGA6: Molecular Evolutionary Genetics Analysis version 6.0. Molecular Biology and Evolution. 2013;30:2725–9.

25. RepeatMasker http://www.repeatmasker.org. Accessed 15 Aug 2015.

26. Zuker M. Mfold web server for nucleic acid folding and hybridization prediction. Nucleic Acids Res. 2003;31(13):3406–15.

27. Telenius H, Carter NP, Bebb CE, Nordenskjold M, Ponder BAJ, Tunnacliffe A. Degenerate oligonucleotide-primed PCR: General amplification of target DNA by a single de-generate primer. Genomics.1992;13:718–25.

28. Galkina S, Deryusheva S, Fillon V, Vignal A, Crooijmans R, Groenen M, Rodionov A, Gaginskaya E. FISH on avian lampbrush chromosomes produces higher resolution gene mapping. Genetica. 2006;128:241–51.

29. Coleman AW. Analysis of mammalian rDNA internal transcribed spacers. PLoS One. 2013;doi: 10.1371/journal.pone.0079122.

30. Bellemain E, Carlsen T, Brochmann C, Coissac E, Taberlet P, Kauserud H. ITS as an environmental DNA barcode for fungi: an in silico approach reveals potential PCR biases. BMC Microbiol. 2010;doi: 10.1186/1471-2180-10-189.

31. Yao H, Song J, Liu C, Luo K, Han J, Li Y, Pang X, Xu H, Zhu Y, Xiao P, Chen S. Use of ITS2 region as the universal DNA barcode for plants and animals. PLoS One. 2010;doi: 10.1371/journal.pone.0013102.

32. Chen S, Yao H, Han J, Liu C, Song J, Shi L, Zhu Y, Ma X, Gao T, Pang X, Luo K, Li Y, Li X, Jia X, Lin Y, Leon C. Validation of the ITS2 region as a novel DNA barcode for identifying medicinal plant species. PLoS One. 2010;doi: 10.1371/journal.pone.0008613.

33. Wang XC, Liu C, Huang L, Bengtsson-Palme J, Chen H, Zhang JH, Cai D, Li JQ. ITS1: a DNA barcode better than ITS2 in eukaryotes? Mol Ecol Resour. 2015;15(3):573–86.

34. Evers R, Grummt I. Molecular coevolution of mammalian ribosomal gene terminator sequences and the transcription termination factor TTF-I. Proc Natl Acad Sci U S A. 1995;92(13):5827–31.

